# Distinct effects of aging and hearing loss on mouse auditory cortex

**DOI:** 10.64898/2026.11.20.768133

**Authors:** Sarah Fouda, Shuzo Sakata

**Affiliations:** Strathclyde Institute of Pharmacy and Biomedical Sciences, University of Strathclyde, 161 Cathedral Street, Glasgow G4 0RE, UK

**Keywords:** aging, auditory cortex, contrast entropy, decoding, spike timing

## Abstract

Age-related hearing loss is accompanied by changes in auditory cortical activity, but it remains unclear which changes reflect aging itself and which depend on the loss of peripheral input. Here we compared spontaneous and sound-evoked activity in the auditory cortex of awake C57BL/6J (C57) mice, which develop early-onset hearing loss, with that of C57BL/6J × CBA/Ca F1 hybrid mice, which retain their hearing. In both strains, aging increased the irregularity of spontaneous firing and reduced the spike-timing precision of narrow-spiking neurons during natural sound presentation. In C57 mice, aging was additionally associated with a greater loss of sound-responsive neurons, increased spontaneous firing of broad-spiking neurons and more variable sound-evoked spike trains. At the level of individual neurons, the effect of sound on contrast entropy, a firing-rate-normalized measure of spike-pattern diversity, also diverged between strains with age and became negative in middle-aged and old C57’s neurons. Nevertheless, population activity discriminated spontaneous, natural sound-evoked and noise-evoked conditions with high accuracy at every age in both strains. Thus, aging and hearing loss have distinct effects on the spiking of individual auditory cortical neurons, whereas population activity continues to discriminate sound conditions.

## Introduction

Age-related hearing loss is among the most common sensory impairments of later life (Lin et al. 2011). It begins in the periphery, with the loss of cochlear hair cells and of the synapses and auditory nerve fibers that carry sound to the brain (Kujawa and Liberman 2009; Sergeyenko et al. 2013; Wu et al. 2019). However, peripheral damage only partly explains the listening difficulties of older adults, and the auditory cortex is itself a site of age-related change (Caspary et al. 2008; Humes et al. 2012; Mittelstadt et al. 2026). It remains poorly characterized how aging and the loss of peripheral input affect auditory cortical processing at the single-neuron level.

Two kinds of change have been described in the aged auditory cortex. First, neuronal responses deteriorate: fewer neurons respond to sound, tuning broadens and temporal precision declines (Turner et al. 2005; de Villers-Sidani et al. 2010; Hughes et al. 2010; Shilling-Scrivo et al. 2021). Second, the loss of peripheral input triggers a compensatory increase in central gain, which raises spontaneous and evoked cortical activity (Kotak et al. 2005; Chambers et al. 2016; Resnik and Polley 2021). Both have been linked to weakened cortical inhibition (Ling et al. 2005; Martin del Campo et al. 2012; Caspary et al. 2013; Ouellet and de Villers-Sidani 2014; Xue et al. 2023). It remains contested whether these changes are driven by peripheral hearing loss or by cortical aging itself. Loss of parvalbumin neurons and of intracortical connectivity occurs in mice that retain their hearing (Rogalla and Hildebrandt 2020; Maximov and Kanold 2025; Xu et al. 2025), which argues for a hearing-independent component.

The C57BL/6J (C57) mouse carries the Cdh23ahl allele and develops progressive, high-frequency-first hearing loss from young adulthood (Willott et al. 1993; Frisina et al. 2011). Crossing C57 animals with CBA/Ca mice, which retain normal hearing into old age, yields F1 hybrids (Hybrids) in which the Cdh23 defect is complemented. These F1 hybrids carry one C57BL/6J and one CBA/Ca genome, and their hearing thresholds remain stable across the lifespan (Lyngholm and Sakata 2019; Bowen et al. 2020). Recording from both strains at young, middle and old ages therefore allows the effects of aging per se, present in both strains, to be distinguished from effects that require peripheral hearing loss, present in C57 mice alone.

Previous descriptions of the aging auditory cortex rest on firing rate and tuning. Here we focus on several quantities to assess neural codes since they depend on the temporal organization of spikes, such as the regularity of spike trains and the precision of spike timing across repeated stimuli (de Ruyter van Steveninck et al. 1997; Schreiber et al. 2003). Additionally, as a complementary measure, we assess contrast entropy, which expresses the entropy of a neuron’s spike patterns relative to the maximum achievable at its firing rate, and so indexes how fully a neuron uses its available patterns independently of how much it fires (Pryluk et al. 2019). Whether aging or hearing loss alters these properties in the auditory cortex, and whether any such changes affect what neuronal populations convey about sound, is unknown.

Here we investigate a previously published dataset (Lyngholm and Sakata 2019), where single-unit activity was recorded from the auditory cortex of awake, head-fixed C57 and Hybrid mice from young adulthood to old age, during silence and during natural sounds.

We report that aging made spontaneous firing more irregular and sound-evoked spike timing less precise in both strains. Several changes were confined to C57 mice: a greater loss of responsive neurons, an age-related rise in spontaneous firing of broad-spiking neurons, more variable sound-evoked spike trains and a sound-induced decrease, rather than increase, in contrast entropy. Despite these differences, population activity discriminated sound conditions accurately at every age in both strains.

## Materials and Methods

### Animals and in vivo electrophysiology

All procedures were performed in accordance with the UK Animals (Scientific Procedures) Act 1986 and were approved by the UK Home Office (PPL 70/8883). Animals, surgery and recording procedures were as described previously (Lyngholm and Sakata 2019). Briefly, we used 89 mice expressing channelrhodopsin-2 in parvalbumin- or somatostatin-positive neurons (PV-Cre or SOM-Cre crossed with Ai32). In C57 mice (35 PV-Cre::Ai32, 18 SOM-Cre::Ai32), both alleles were on the C57BL/6J background. In Hybrid mice (19 PV-Cre::Ai32, 17 SOM-Cre::Ai32), the Ai32 allele was on the CBA/Ca background, giving C57BL/6J × CBA/Ca F1 hybrids in which the Cdh23 defect of C57 mice is complemented and hearing is preserved (Lyngholm and Sakata 2019). Optogenetic stimulation was not analyzed in this study. Mice were grouped by age at recording as young (< 190 days), middle-aged (190– 454 days) or old (≥ 455 days) (Flurkey et al. 2007).

Mice were implanted with a head-post and habituated to head fixation. Recordings were made in awake, head-fixed mice from the primary auditory cortex with 32- or 64-channel silicon probes (NeuroNexus), and signals were digitized at 20 kHz (RHD2000, Intan Technologies).

### Sound stimulation

Recordings were performed inside a sound-attenuating chamber lined with acoustic foam (MAC-3, IAC Acoustics). Auditory stimuli were generated digitally at 97.7 kHz with an RZ6 processor (Tucker-Davis Technologies) and delivered through a calibrated ES1 electrostatic speaker (Tucker-Davis Technologies) placed ∼15 cm in front of the animal.

A silent period of ∼20 min was recorded first to characterize spontaneous activity, followed by approximately 15 min of natural sound stimulation. Natural stimuli were 10-s broadband environmental sound sequences, presented at 65 dB sound pressure level (SPL), with 30–100 repetitions depending on the recording and a 1-s inter-stimulus interval.

In 33 mice (16 C57, 17 Hybrid), 100-ms white-noise bursts were additionally presented at intensities from 10 to 90 dB SPL in 10-dB steps, with 15 to 100 repetitions per intensity and a 300-ms inter-stimulus interval.

### Spike sorting and cell-type classification

Spike sorting was performed as described previously (Lyngholm and Sakata 2019). Only well-isolated single units with an isolation distance greater than 20 were retained (Schmitzer-Torbert et al. 2005).

For each isolated unit, the trough-to-peak duration of the mean spike waveform was measured. Units with a trough-to-peak duration above 0.55 ms were classified as broad-spiking (BS, putative excitatory neurons) and units at or below 0.55 ms as narrow-spiking (NS, putative fast-spiking inhibitory neurons) (**Fig. 1E**) (Lyngholm and Sakata 2019).

**Figure 1.**
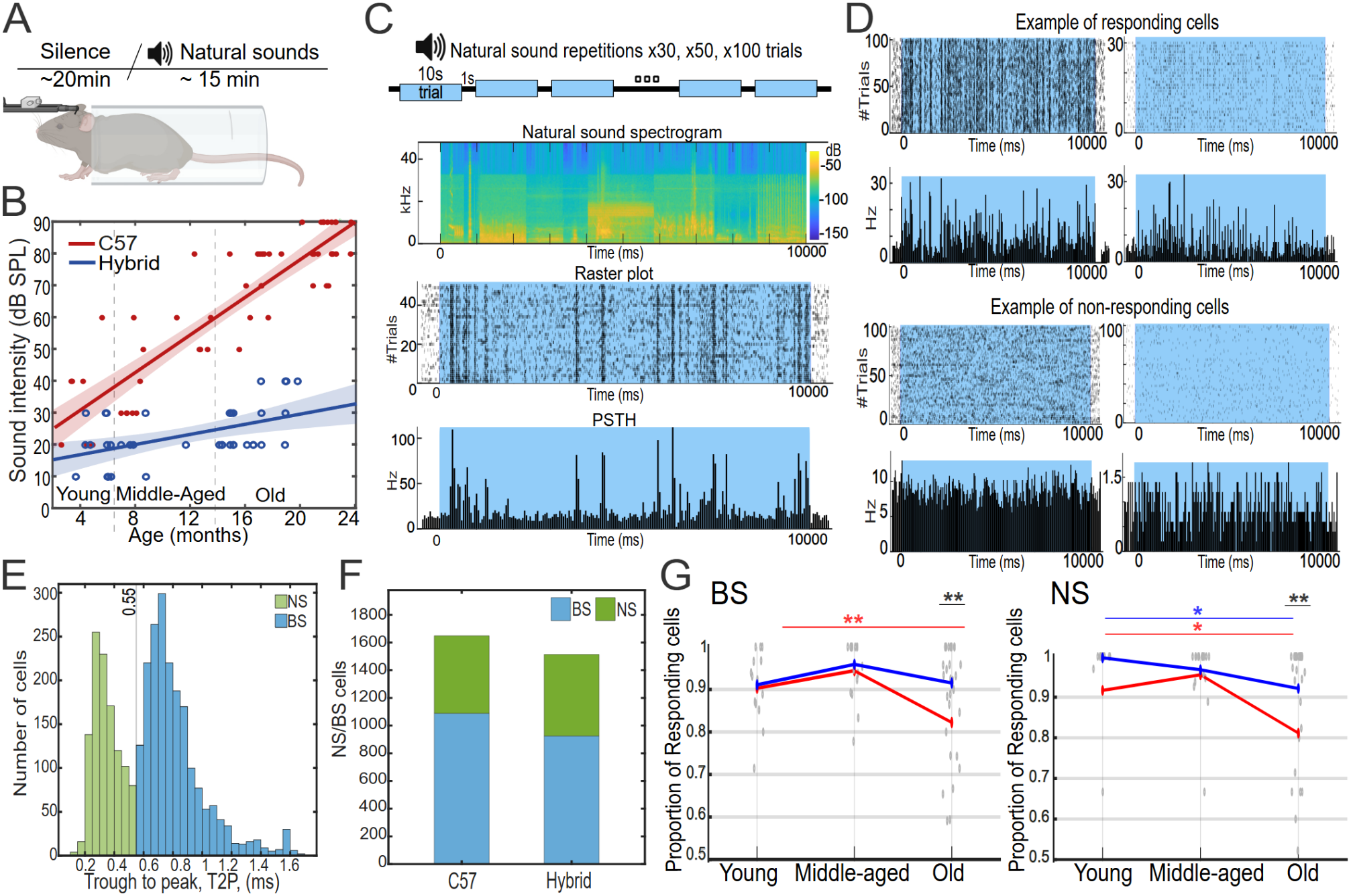
Age-dependent alterations in auditory responsiveness in C57 and Hybrid mice. **(A)** Experimental paradigm for natural sound stimulation. Following a silent period of approximately 20 min, mice were exposed to repeated presentations of natural sounds for approximately 15 min. **(B)** Hearing thresholds (dB SPL) as a function of age, re-plotted from Lyngholm and Sakata (2019). The schematic indicates the Cdh23 mutation associated with hearing loss in C57 mice and its restoration in the Hybrid background. **(C)** Neuronal responses to repeated natural sound presentations. The natural sound spectrogram is shown together with trial-aligned raster plots and peri-stimulus time histograms (PSTHs), illustrating stimulus-locked neuronal activity. **(D)** Representative examples of responsive and non-responsive neurons. Neurons were classified as responsive when evoked firing exceeded the pre-stimulus baseline firing (250 ms before stimulus onset) by 3 SD. **(E)** Distribution of trough-to-peak duration for recorded neurons, separated into broad-spiking (BS) and narrow-spiking (NS) populations. The dashed line at 0.55 ms indicates the boundary used to distinguish the two populations. **(F)** Number of BS and NS neurons recorded in C57 and Hybrid mice. Bars show all recorded neurons (C57, n = 1,687: 1,114 BS, 573 NS; Hybrid, n = 1,514: 925 BS, 589 NS). **(G)** Proportion of BS and NS responsive cells across young, middle-aged, and old age groups in C57 and Hybrid mice, computed on the natural-sound dataset (2,179 neurons). Gray dots represent individual recordings; colored lines indicate the mean proportion for each strain (red, C57; blue, Hybrid). Significant differences were observed between strains in the old group for both BS (p = 0.000168) and NS cells (p = 0.00295). Within C57 mice, the proportion of responsive cells decreased from young to old for both BS (p = 0.00204) and NS cells (p = 0.0307). Within Hybrid mice, a significant young-to-old difference was observed for NS cells (p = 0.0115). Only significant comparisons are indicated.

### Sound responsiveness

Spike trains were binned at 50 ms. For natural sounds, the baseline firing rate and its standard deviation were computed over the 250 ms preceding stimulus onset, and the evoked window comprised the 10-s stimulus plus the 250 ms following stimulus offset. A neuron was classified as responsive when

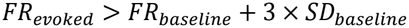

and as non-responsive otherwise, where FR denotes firing rate and SD the standard deviation.

### Firing rate and spike-train variability

Firing rate was computed as the number of spikes divided by the duration of the analysis window. Neurons with a mean firing rate below 0.1 Hz were excluded. Spike-train variability was quantified as the coefficient of variation (CV) of inter-spike intervals (ISIs):

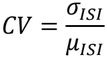

where σ and μ are the standard deviation and the mean of the ISIs. For spontaneous activity, ISIs were computed over the silent period. For sound-evoked activity, ISIs were computed within each natural sound presentation and pooled across presentations.

### Spike-timing precision

The precision of spike timing across repeated natural sounds was quantified with the Schreiber similarity index (Schreiber et al. 2003). Spike trains from individual repetitions were convolved with a Gaussian kernel, and the index was computed as the mean normalized inner product over all pairs of repetitions:

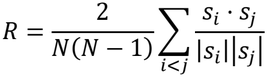

where s denotes the smoothed spike train of a repetition and N the number of repetitions. The index ranges from 0 (no reproducible temporal structure) to 1 (identical responses). Kernel widths (σ) of 1, 4, 16 and 32 ms were evaluated, and σ = 16 ms was used for all analyses.

### Contrast entropy

Contrast entropy (CE) quantifies how much of the available spike-pattern space a neuron uses, relative to a random spike train of the same firing rate (Pryluk et al. 2019). Two parameters define the analysis. The letter bin width (Δt) is the bin size used to convert a spike train into a binary sequence, in which a bin is 1 if it contains at least one spike and 0 otherwise. The word length (W) is the number of consecutive letters that form a word. The binary sequence was divided into non-overlapping words, and the probability of each word was estimated from its frequency. The empirical entropy rate (ER) is

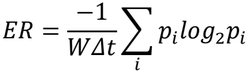

where p with subscript i is the probability of word i (Rieke et al. 1999). The analytic entropy rate (EA) is the entropy rate expected for a neuron with the same firing rate (FR) whose spikes occur independently in each bin. The probability that a bin contains a spike is

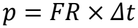

and

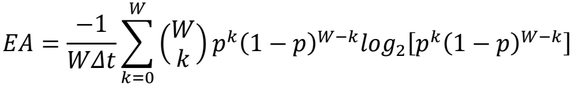

where Δt is expressed in seconds and k is the number of spike-containing bins in a word. CE is the ratio of the two:

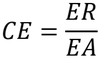

We computed CE for bin widths of 1, 2, 4, 8 and 16 ms and word lengths of 4, 8 and 16 letters (**Fig. S1C**). All reported analyses use Δt = 4 ms and W = 8, giving 32-ms words and 256 possible words. Neurons with a mean firing rate below 0.1 Hz were excluded (62 of 2,179 neurons in the natural-sound dataset).

To quantify the effect of sound within a neuron, we computed the CE modulation index for the 1,735 neurons with both spontaneous and sound-evoked estimates:

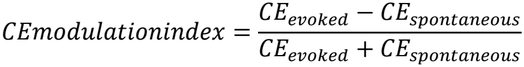

### Population decoding

We tested whether population activity discriminates three conditions: spontaneous, natural sound-evoked and noise-evoked activity. The analysis was restricted to the 33 mice in which all three conditions were recorded. Spike counts were computed in 100-ms windows, which matches the duration of the noise bursts so that the three conditions were compared over equal durations. Windows were aligned to the onset of natural sound and noise presentations, and spontaneous activity was divided into consecutive 100-ms windows within the silent period. The three conditions were balanced within each mouse by randomly drawing the same number of windows from each, up to 100 per condition. Noise windows were pooled across intensities without balancing across intensities. Decoding was run for all neurons and separately for BS and NS neurons.

We used a template-correlation decoder with leave-one-out cross-validation. For each condition k, the template was the mean population vector of the training windows of that condition, and each held-out window was assigned to the condition whose template it correlated with most strongly:

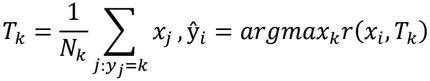

where x denotes the population vector of a window, T the template and r the Pearson correlation coefficient. Accuracy was the proportion of correctly classified windows, with a chance level of 33.3%, and errors were summarized in a confusion matrix. In the main analysis (**Fig. 5**), decoding was applied to spike counts from each recording and summarized per mouse. To test whether the results depended on the decoder, we repeated the analysis with a linear support vector machine and linear discriminant analysis on the same windows, after concatenating all neurons of a mouse and z-scoring spike counts within each cross-validation fold using the training windows only (**Fig. S2**).

### Statistical analysis

Analyses were performed in MATLAB (MathWorks) and R. The neuron was the unit of analysis for single-neuron measures and the mouse for population decoding. Proportions of responsive neurons were compared with two-sample tests for equality of proportions with Yates’ continuity correction. Two groups were compared with two-sided Wilcoxon rank-sum tests, and paired comparisons within neurons with Wilcoxon signed-rank tests. Associations were assessed with Spearman rank correlation or linear regression. Effects of strain and age were tested with two-way permutation ANOVA (9,999 permutations) (Anderson 2001), followed by planned post hoc permutation tests (999 permutations) with Holm correction (Holm 1979); p values were obtained from the permutation distributions. F statistics are reported with their degrees of freedom (strain, 1; age, 2; strain × age, 2; error, number of neurons minus 6). Decoding accuracy was compared with chance using one-sided one-sample Wilcoxon signed-rank tests, across strains and age groups with two-way permutation ANOVA (error degrees of freedom, number of mice minus 6) and Kruskal–Wallis tests, and against age with Spearman rank correlation. Decoders were compared with paired t-tests with Holm correction (**Fig. S2**). Significance was set at p < 0.05.

### Use of AI tools

Analysis scripts were optimized using ChatGPT. All AI-generated code was reviewed, validated, and revised by the authors, who confirmed accuracy and reproducibility of results.

During the preparation of the manuscript, the authors used Claude and ChatGPT in order to help summarize background literature notes, check statistical information and code, and proofread the manuscript. After using these tools, the authors reviewed, verified, and edited the content as needed and take full responsibility for the accuracy of the manuscript.

## Results

We recorded single-unit activity from the auditory cortex of 89 awake head-fixed mice of two strains across age. C57BL/6J (C57) mice carry a *Cdh23*^*ahl*^ mutation and develop early-onset hearing loss, whereas F1 hybrid of C57 and CBA/Ca (Hybrid) mice carry a restored *Cdh23* allele and retain near-normal hearing into old age (Lyngholm and Sakata 2019). Mice were assigned to three age groups (Flurkey et al. 2007): young (< 190 days), middle-aged (190 to 454 days) and old (≥ 455 days). Each recording session began with approximately 20 min of silence and continued with approximately 15 min of repeated 10-s natural sounds (**Fig. 1A**). Neurons were classified into broad-spiking (BS) and narrow-spiking (NS) populations based on their trough-to-peak duration of the average spike waveform (**Fig. 1E**). The recorded sample comprised 1,687 neurons from C57 mice (1,114 BS and 573 NS) and 1,514 neurons from Hybrid mice (925 BS and 589 NS) (**Fig. 1F**).

### Age-related loss of auditory responsiveness is stronger in C57 than in Hybrid mice

We first asked whether the proportion of sound-responsive cortical neurons declines with age, and whether peripheral hearing loss amplifies that decline. Hearing thresholds rose with age in C57 mice but remained stable in Hybrid mice, consistent with the presence of the *Cdh23* mutation in the former and its restoration in the latter (**Fig. 1B**); thresholds were re-plotted from Lyngholm and Sakata (2019). Repeated presentations of the same natural sound evoked stimulus-locked firing (**Fig. 1C**). Neurons were classified as responsive when evoked firing exceeded baseline firing by 3 SD, and as non-responsive otherwise (**Fig. 1D**). The proportion of responsive neurons declined from young to old age in C57 mice for both cell classes, and in Hybrid mice for NS neurons only (**Fig. 1G**). Within C57 mice, the proportion of responsive neurons fell from young to old age in BS neurons (p = 0.00204) and in NS neurons (p = 0.0307). Within Hybrid mice, the same young-to-old comparison was significant only for NS neurons (p = 0.0115). In old mice the two strains differed for both cell classes, with fewer responsive neurons in C57 than in Hybrid mice among BS neurons (p = 0.000168) and NS neurons (p = 0.00295). These results indicate that aging reduces the fraction of auditory cortical neurons that respond to natural sounds in C57 mice, where the reduction affected both BS and NS populations and left old C57 mice with fewer responsive neurons than old Hybrid mice.

### Spontaneous and sound-evoked firing change with age in a strain- and cell-type-specific manner

We next asked whether aged auditory cortex simply fires less, or whether spontaneous and sound-evoked activity follow separate trajectories across three age groups. We found that spontaneous firing rate diverged between the two strains (**Fig. 2A**). In C57’s BS neurons, spontaneous firing increased from young to old age (p = 0.04). In Hybrid’s BS neurons, it decreased from middle age to old age (p = 0.024) and from young to old age (p = 0.018). The two strains consequently differed in young and in old mice (both p = 0.018). In NS neurons, spontaneous firing decreased significantly with age in Hybrid mice (p = 0.009), whereas no significant age-related change was observed in C57 mice. A significant difference between strains was also observed in the old group (p = 0.009). Sound-evoked firing followed a different pattern. Evoked firing rate of BS neurons did not change significantly with age in either strain, whereas evoked firing of NS neurons showed a main effect of age (F(2, 799) = 11.40, p = 0.0001) and declined with age in C57 mice (young to old, p = 0.049) and in Hybrid mice (young to middle age, p = 0.009; young to old, p = 0.009) (**Fig. 2A**).

**Figure 2.**
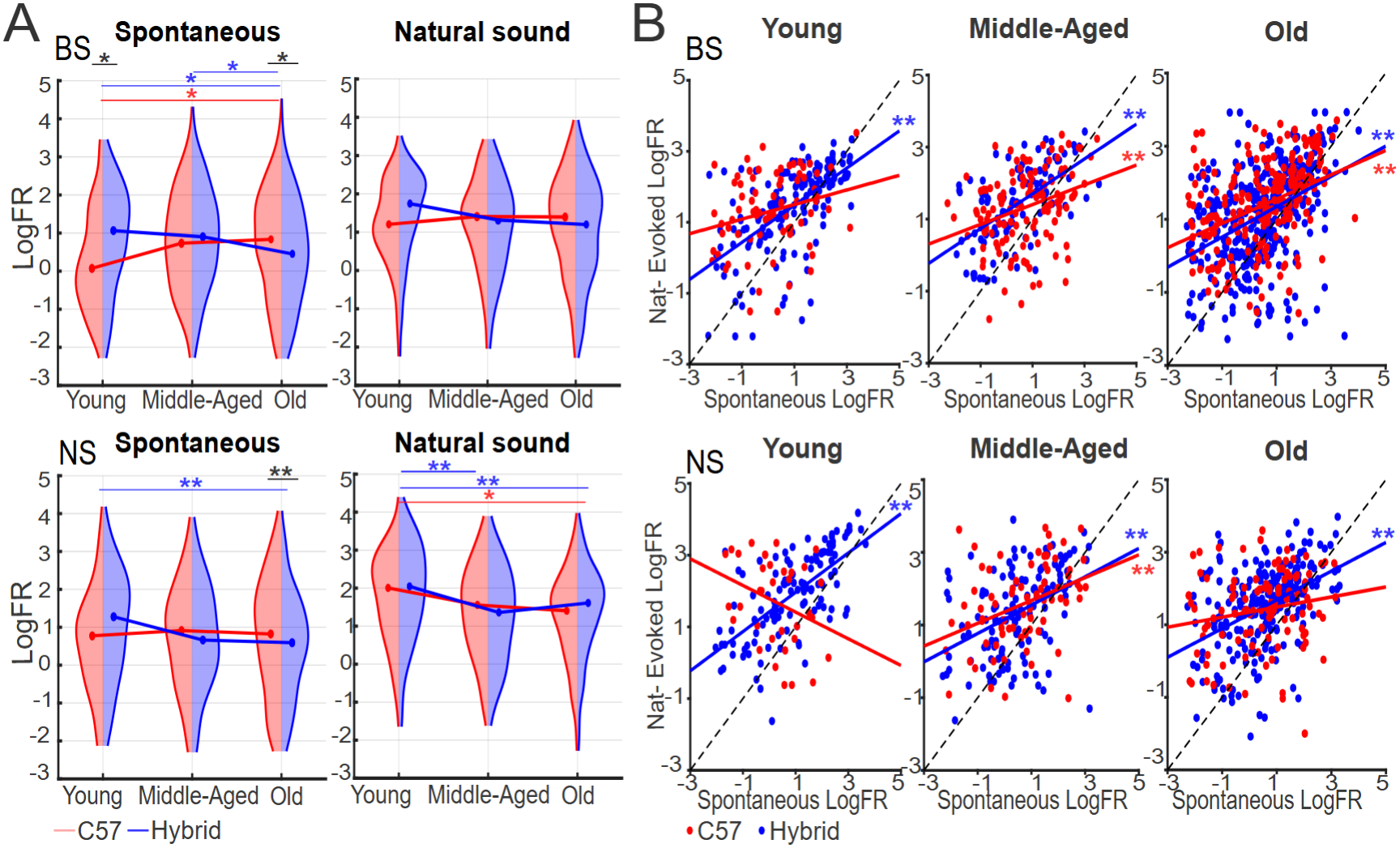
Age-related changes in spontaneous and natural sound-evoked firing rates. **(A)** Distributions of spontaneous and natural sound-evoked log-transformed firing rate (LogFR) in BS (top) and NS (bottom) neurons across age groups in C57 (red) and Hybrid (blue) mice. Horizontal lines indicate medians. For spontaneous activity, significant differences were detected in BS between C57 young and old (p = 0.04), Hybrid young and old (p = 0.018), Hybrid middle-aged and old (p = 0.024), and C57 versus Hybrid in young (p = 0.018) and old (p = 0.018) mice. In NS, significant differences were observed between young and old Hybrid mice (p = 0.009) and between C57 and Hybrid in old group (p = 0.009). For natural sound-evoked activity, significant differences were detected in NS between C57 young and old (p = 0.049), Hybrid young and middle-aged (p = 0.009), and Hybrid young and old (p = 0.009). **(B)** Relationship between spontaneous and natural sound-evoked LogFR across young, middle-aged and old C57 and Hybrid mice. Each point represents an individual neuron; solid lines indicate linear regressions. For BS neurons, spontaneous and sound-evoked LogFR were positively associated in all Hybrid groups: Hybrid young (β = 0.523, R^2^ = 0.346, p = 6.97 × 10^−16^), Hybrid middle-aged (β = 0.483, R^2^ = 0.342, p = 5.43 × 10^−11^), and Hybrid old (β = 0.414, R^2^ = 0.175, p = 1.36 × 10^−16^), and in the C57 middle-aged and old groups: C57 middle-aged (β = 0.274, R^2^ = 0.102, p = 2.61 × 10^−4^), C57 old (β = 0.332, R^2^ = 0.156, p = 7.90 × 10^−10^); not in C57 young (β = 0.169, R^2^ = 0.040, p = 0.094). For NS neurons, significant positive associations were observed in Hybrid young (β = 0.548, R^2^ = 0.429, p = 1.44 × 10^−15^), Hybrid middle-aged (β = 0.390, R^2^ = 0.173, p = 8.57 × 10^−7^), C57 middle-aged (β = 0.314, R^2^ = 0.136, p = 0.001), and Hybrid old (β = 0.395, R^2^ = 0.200, p = 9.83 × 10^−13^); associations were not significant in C57 young (β = −0.434, R^2^ = 0.122, p = 0.0504) or C57 old (β = 0.139, R^2^ = 0.032, p = 0.0624).

We then asked whether spontaneous activity predicted the magnitude of sound-evoked responses (**Fig. 2B**). In Hybrid BS neurons, spontaneous and evoked firing rates were positively associated at every age (Hybrid young, β = 0.523, R^2^ = 0.346, p = 6.97 × 10^−16^; middle-aged, β = 0.483, R^2^ = 0.342, p = 5.43 × 10^−11^; old, β = 0.414, R^2^ = 0.175, p = 1.36 × 10^−16^). In C57 BS neurons, significant positive associations were observed in middle-aged and old mice, but not in young mice (young, β = 0.169, R^2^ = 0.040, p = 0.094; middle-aged, β = 0.274, R^2^ = 0.102, p = 2.61 × 10^−4^; old, β = 0.332, R^2^ = 0.156, p = 7.90 × 10^−10^). The slopes were numerically shallower in C57 than in Hybrid mice across all age groups.

In NS neurons, the association was significant in Hybrid mice at every age (Hybrid young, β = 0.548, R^2^ = 0.429, p = 1.44 × 10^−15^; middle-aged, β = 0.390, R^2^ = 0.173, p = 8.57 × 10^−7^; old, β = 0.395, R^2^ = 0.200, p = 9.83 × 10^−13^). In C57 mice, the association was significant only in middle-aged animals (β = 0.314, R^2^ = 0.136, p = 0.001). No significant association was observed in young C57 mice (β = −0.434, R^2^ = 0.122, p = 0.0504) or old C57 mice (β = 0.139, R^2^ = 0.032, p = 0.0624). These results indicate that aging lowers spontaneous auditory cortical firing in Hybrid mice, whereas hearing loss in C57 mice is accompanied by an age-related rise in spontaneous firing of BS neurons and by weaker coupling between spontaneous and sound-evoked activity.

### Spike timing changes with age in a strain- and cell-type-specific manner

Given these changes in firing rate, we asked whether the temporal structure of spike trains also changes with age. The irregularity of spontaneous firing, quantified as the coefficient of variation (CV) of inter-spike intervals, increased with age in both strains and both cell classes (**Fig. 3A**). CV increased from young to middle-aged (p = 5.6 × 10^−7^) and from young to old (p = 0.007) in C57 BS neurons, and from young to old in Hybrid BS neurons (p = 2.4 × 10^−4^). In NS neurons, CV increased from young to middle-aged (p = 0.02) in C57 mice, and from young to old (p = 9.0 × 10^−5^) and from middle-aged to old (p = 7.6 × 10^−4^) in Hybrid mice. During repeated natural sound presentations, responsive cells defined in **Fig. 1** exhibited separate trajectories in the two strains (**Fig. 3B**): In C57 mice, CV increased from young to old age in BS neurons (p = 0.0008) and from young to middle-aged and young to old in NS neurons (both p = 0.0006). In Hybrid mice, CV instead fell from middle-aged to old age in BS neurons (median 1.340 to 1.202, p = 0.0021) and followed an inverted U-shaped trajectory in NS neurons (young to middleaged, p = 0.029; middle-aged to old age, p = 0.024). As a result, variability during natural sounds was higher in C57 than in Hybrid mice in old BS neurons (p = 0.0008) and in middle-aged and old NS neurons (both p = 0.0006).

**Figure 3.**
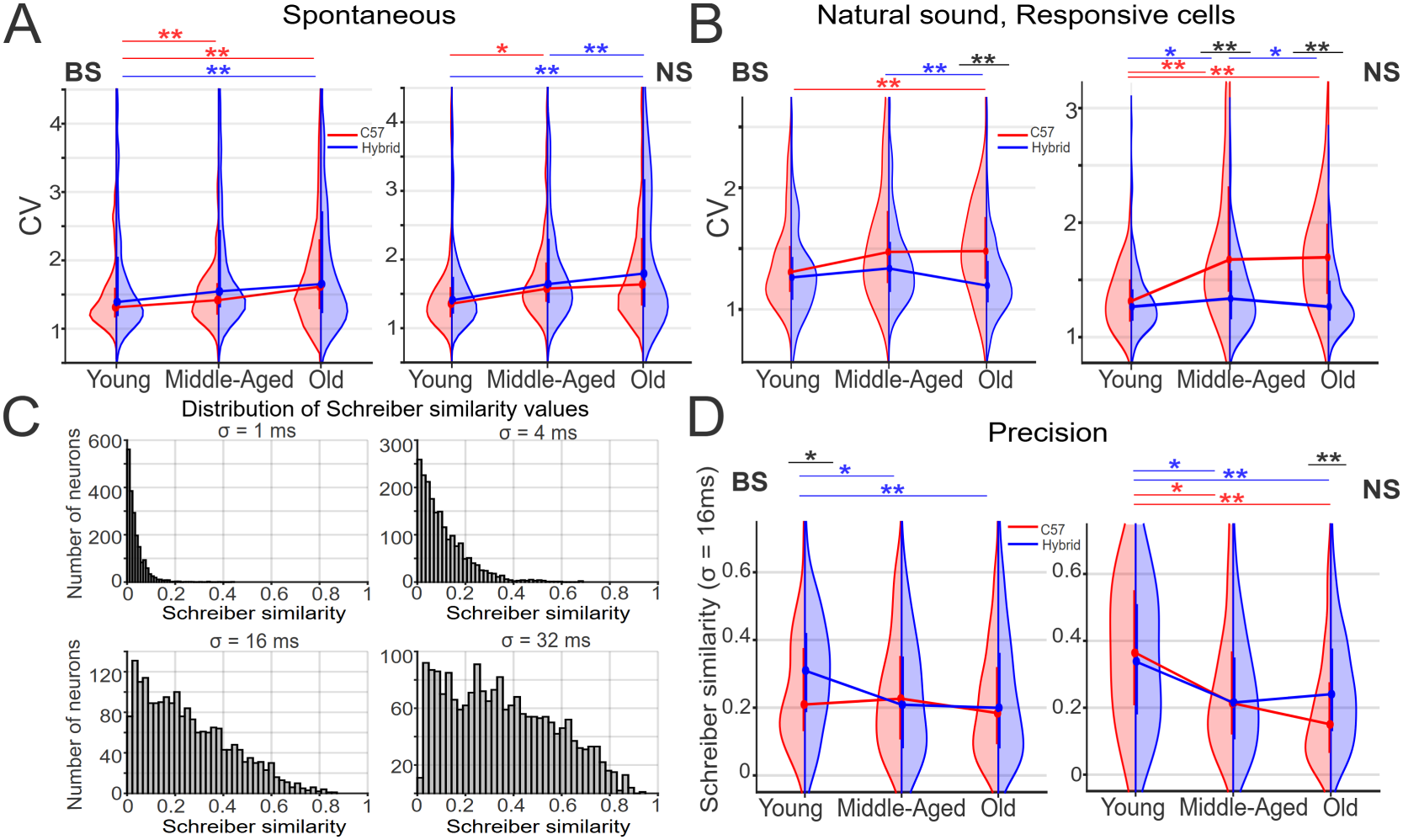
Age-related changes in spike-train variability and spike-timing precision. **(A)** Coefficient of variation (CV) of spontaneous firing in BS (left) and NS (right) neurons from C57 (red) and Hybrid (blue) mice across age. CV increased in C57 BS neurons from young to middle-aged (p = 5.6 × 10^−7^) and old (p = 0.007), and in Hybrid BS neurons from young to old (p = 2.4 × 10^−4^). In NS neurons, CV increased in C57 mice from young to middleaged (p = 0.02), and in Hybrid mice from young to old (p = 9.0 × 10^−5^) and from middle-aged to old (p = 7.6 × 10^−4^). Half-violins show individual-neuron distributions; circles indicate medians, vertical bars indicate IQR, and lines connect agegroup medians. **(B)** CV of sound-evoked responses during repeated natural sound presentations in BS (left) and NS (right) neurons. In BS neurons, CV increased from young to old mice in C57 mice (p = 0.0008), whereas it decreased from middle-aged to old mice in Hybrid mice (p = 0.0021). In NS neurons, CV increased from young to middle-aged (p = 0.0006), and from young to old mice in C57 mice (p = 0.0006). In Hybrid mice, it increased from young to middle-aged mice (p = 0.029) and then decreased from middle-aged to old mice (p = 0.024). CV was higher in C57 than in Hybrid mice in old BS neurons (p = 0.0008) and in middle-aged and old NS neurons (both p = 0.0006). **(C)** Distribution of Schreiber similarity values computed across repeated natural sound presentations using Gaussian smoothing windows of σ = 1, 4, 16, and 32 ms. Increasing the smoothing window shifts the distribution toward higher similarity values. A smoothing window of σ = 16 ms was selected for subsequent analyses as a compromise between temporal precision and robustness to spike-time variability. **(D)** Spike-timing precision was computed as Schreiber similarity index (σ = 16 ms) (Schreiber et al. 2003) of repeated natural sound responses in responsive BS (left) and NS (right) neurons. In BS neurons, precision remained stable across age groups in C57 mice but decreased in Hybrid mice from young to middle-aged and from young to old (p = 0.011 and p = 0.0009, respectively), with higher precision in young Hybrid than in young C57 mice (p = 0.018). In NS neurons, precision decreased from young to middle-aged and from young to old in both C57 (p = 0.014 and p = 0.0009, respectively) and Hybrid mice (p = 0.013 and p = 0.0021, respectively), with higher precision in old Hybrid than in old C57 mice (p = 0.0016).

We also measured the precision of spike timing across repeated presentations using the Schreiber similarity index (Schreiber et al. 2003). The index was computed for Gaussian kernels of σ = 1, 4, 16 and 32 ms (**Fig. 3C**), and σ = 16 ms was retained for all further analyses as a compromise between temporal precision and robustness to spike-time jitter. We found that spike-timing precision declined with age (**Fig. 3D**). In BS neurons, precision remained stable across age groups in C57 mice but decreased in Hybrid mice from young to middle-aged and from young to old (p = 0.011 and p = 0.0009, respectively), with higher precision in young Hybrid mice than in young C57 mice (p = 0.018). In NS neurons, precision decreased from young to middle-aged and from young to old in both C57 (p = 0.014 and p = 0.0009, respectively) and Hybrid mice (p = 0.013 and p = 0.0021, respectively). Precision was also higher in old Hybrid mice than in old C57 mice (p = 0.0016). These results suggest that aging impairs the temporal precision of sound-evoked responses, with a greater age-related loss of precision in C57 than in Hybrid mice for NS neurons.

### Contrast entropy diverges between strains with age

The findings above raised the question of whether the age- and strain-dependent changes in auditory cortical processing were accompanied by changes in the temporal structure of spike trains that firing rate alone cannot explain. To address this, we further computed contrast entropy (CE), the empirical entropy rate of a binarized spike train divided by the analytic entropy expected for a Poisson-like neuron with the same firing rate (Pryluk et al. 2019). This normalization expresses the entropy of the spike train relative to its theoretical maximum and therefore measures how much of the available spike-pattern space a neuron uses, independently of how fast it fires.

We first verified that CE behaves as intended (**Fig. S1**). We computed both analytical and empirical entropy rates in each neuron given its firing rate (**Fig. S1A**). The empirical entropy rate increased steeply with firing rate (**Fig. S1A**), whereas the conversion to CE removed that dependence during both spontaneous activity (from r = 0.97 to r = −0.15) and natural sound presentation (from r = 0.96 to r = −0.17) (**Fig. S1B**), indicating that CE is independent of firing rate.

CE relies on two key variables: letter bin width and word length. The former is the bin size to binarize a spike train, whereas the latter is the size of letter sequences. We systematically swept letter bin widths (1 to 16 ms) and word lengths (4, 8, 16 letters) to compute CE (**Fig. S1C**). As expected, the median of CE decreases in longer bins and words due to sample size and differences in correlation between successive words. Examples of the difference between the entropy rate of neurons and the optimal word distributions, computed with 4 letters per word, are shown in **Fig. S1D**. In the following analysis, we used a bin width of 4 ms with 8 letters per word. We also confirmed that CE estimates from the first and second halves of each recording agreed closely (**Fig. S1E**).

We then examined how CE changes with age (**Fig. 4**). During spontaneous activity, CE in BS neurons showed effects of strain (F(1, 1789) = 5.04, p = 0.0018) and age (F(2, 1789) = 3.02, p = 0.0146) without an interaction (F(2, 1789) = 2.61, p = 0.072) (**Fig. 4A**). CE in Hybrid’s BS neurons decreased from young to middle age (p = 0.009), and C57’s BS neurons had higher spontaneous CE than Hybrid’s BS neurons at middle age (p = 0.016). Spontaneous CE in NS neurons showed no effect of strain (p = 0.18), age (p = 0.22) or their interaction (p = 0.68).

**Figure 4.**
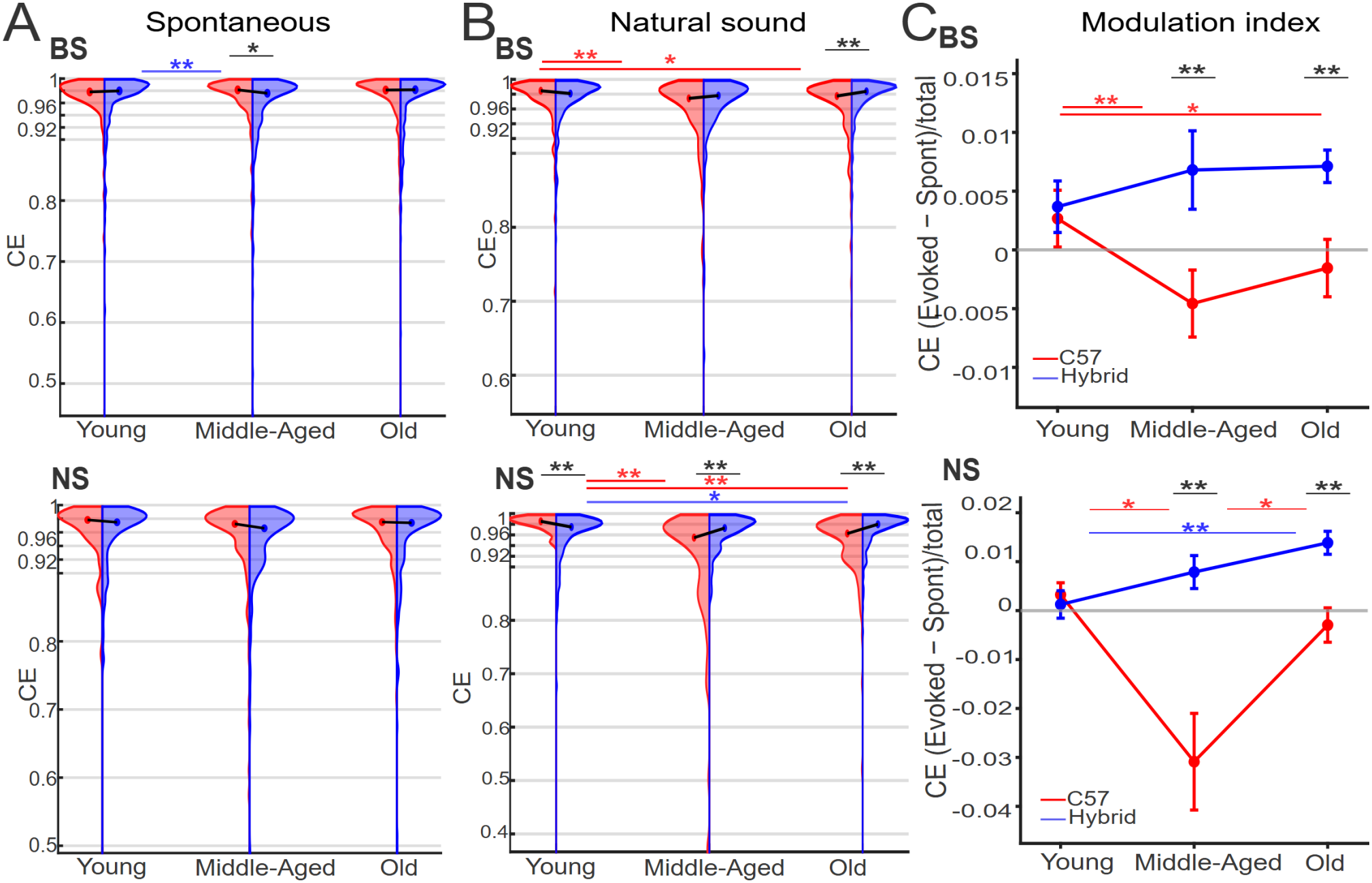
Age- and strain-dependent changes in contrast entropy during spontaneous and natural sound-evoked activity. **(A)** Age-dependent changes in spontaneous CE. In BS cells, a significant strain effect (F = 5.04, p = 0.0018) and age effect (F = 3.02, p = 0.0146) were observed (strain × age interaction not significant, p = 0.072): CE decreased from young to middle-aged in Hybrid mice (p = 0.009) and was lower in Hybrid than C57 mice at middle age (p = 0.016). In NS cells, no significant effects of strain, age, or their interaction were detected (all p ≥ 0.18). **(B)** Age-dependent changes in CE during natural sound presentation. In BS cells, significant effects of strain (F = 8.02, p = 0.0001), age (F = 4.19, p = 0.0027), and strain × age interaction (F = 4.11, p = 0.0158) were observed: CE decreased from young to middle-aged (p = 0.009) and young to old (p = 0.014) in C57 mice, and was lower in C57 than Hybrid mice in old animals (p = 0.009). In NS cells, significant effects of strain (F = 29.67, p = 0.0001), age (F = 15.71, p = 0.0001), and strain × age interaction (F = 22.75, p = 0.0001) were observed: CE decreased from young to middle-aged and young to old in C57 mice (both p = 0.009) and increased from young to old in Hybrid mice (p = 0.048); CE was higher in C57 than Hybrid mice in young animals but lower in middle-aged and old animals (all p = 0.009). **(C)** CE modulation index, (CE_evoked − CE_spontaneous)/(CE_evoked + CE_spontaneous), across age groups in BS (left) and NS (right) cells (matched cells, n = 1,735). In BS cells, a significant strain effect was observed (F = 6.54, p = 0.0004; age and strain × age interaction not significant, p = 0.39 and p = 0.18, respectively): the index decreased from young to middle-aged (p = 0.009) and young to old (p = 0.018) in C57 mice, and was lower in C57 than Hybrid mice at middle age (p = 0.009) and old age (p = 0.009). In NS cells, significant effects of strain (F = 16.53, p = 0.0001), age (F = 7.46, p = 0.0001), and strain × age interaction (F = 7.93, p = 0.0003) were observed: the index decreased from young to middle-aged in C57 mice (p = 0.018), increased from middle-aged to old in C57 mice (p = 0.035), increased from young to old in Hybrid mice (p = 0.009), and was lower in C57 than Hybrid mice at middle age (p = 0.009) and old age (p = 0.009).

During natural sound presentation, the two strains diverged with age (**Fig. 4B**). In BS neurons, there were effects of strain (F(1, 1306) = 8.02, p = 0.0001), age (F(2, 1306) = 4.19, p = 0.0027) and their interaction (F(2, 1306) = 4.11, p = 0.0158): CE in C57 mice fell from young to middle age (p = 0.009) and from young to old age (p = 0.014), and was lower than in Hybrid mice at old age (p = 0.009). In NS neurons, the interaction was larger (strain, F(1, 799) = 29.67, p = 0.0001; age, F(2, 799) = 15.71, p = 0.0001; strain × age, F(2, 799) = 22.75, p = 0.0001). CE in C57’s NS neurons fell from young to middle age and from young to old age (both p = 0.009), whereas in Hybrid’s NS neurons it rose from young to old age (p = 0.048). The strains differed at every age (all p = 0.009), with young C57’s NS neurons showing higher evoked CE than Hybrid’s NS neurons and middleaged and old C57’s NS neurons showing lower evoked CE.

To compare the two conditions within the same neurons, we computed a CE modulation index for the 1,735 neurons recorded in both conditions (**Fig. 4C**). In BS neurons, the index showed a strain effect (F(1, 1037) = 6.54, p = 0.0004) with no age effect (p = 0.39) or interaction (p = 0.18): it decreased in C57 mice from young to middle age (p = 0.009) and from young to old age (p = 0.018), and was lower in C57 than Hybrid mice at middle age (p = 0.009) and old age (p = 0.009). In NS neurons, all three effects were significant (strain, F(1, 686) = 16.53, p = 0.0001; age, F(2, 686) = 7.46, p = 0.0001; strain × age, F(2, 686) = 7.93, p = 0.0003): the index decreased in C57 mice from young to middle age (p = 0.018), increased from middle age to old age (p = 0.035), increased in Hybrid mice from young to old age (p = 0.009), and was lower in C57 than Hybrid mice at middle and old age (both p = 0.009). Median values were positive in Hybrid’s neurons at all ages but negative in middle-aged and old C57’s neurons. Thus, at the level of individual neurons, the effect of sound presentations on CE diverged between strains with age: it was slightly positive in Hybrid’s neurons and became negative in middle-aged and old C57’s neurons.

### Population activity discriminates sound conditions at every age in both strains

Finally, we asked whether these age- and strain-dependent changes in single neurons affect how well population activity distinguishes sound conditions. In 33 mice (16 C57 and 17 Hybrid), sessions included 100-ms white-noise bursts in addition to silence and natural sounds. For each mouse, population vectors of spike counts in 100-ms windows were assigned to one of three conditions (spontaneous, natural sound-evoked or noise-evoked) with a template-correlation decoder under leave-one-out cross-validation (see Materials and Methods). Decoding was performed for all neurons and separately for BS and NS neurons.

Decoding accuracy was well above the chance level of 33.3% at every age in both strains. Across all neurons, each condition was classified above chance (spontaneous, 73.66 ± 16.31%; natural sound, 91.39 ± 7.06%; noise, 71.36 ± 19.62%; mean ± SD; all p < 10^−^6, one-sided Wilcoxon signed-rank tests against chance) and accuracy did not differ between strains (one-way permutation ANOVA with Holm correction, strain effect: all neurons, p = 0.2118; BS, p = 0.2118; NS, p = 0.3247).

Overall accuracy was 79.62 ± 9.09% in young, 75.38 ± 10.26% in middle-aged and 81.96 ± 8.56% in old mice and did not differ among age groups (Kruskal– Wallis test, p = 0.246). The same held for BS (p = 0.215) and NS populations (p = 0.997). Consistent with these findings, two-way permutation ANOVAs revealed no significant main effects of strain or age group, or strain × age interactions, in the All, BS or NS populations (all p-values ≥ 0.120; Fig. 5A).

**Figure 5.**
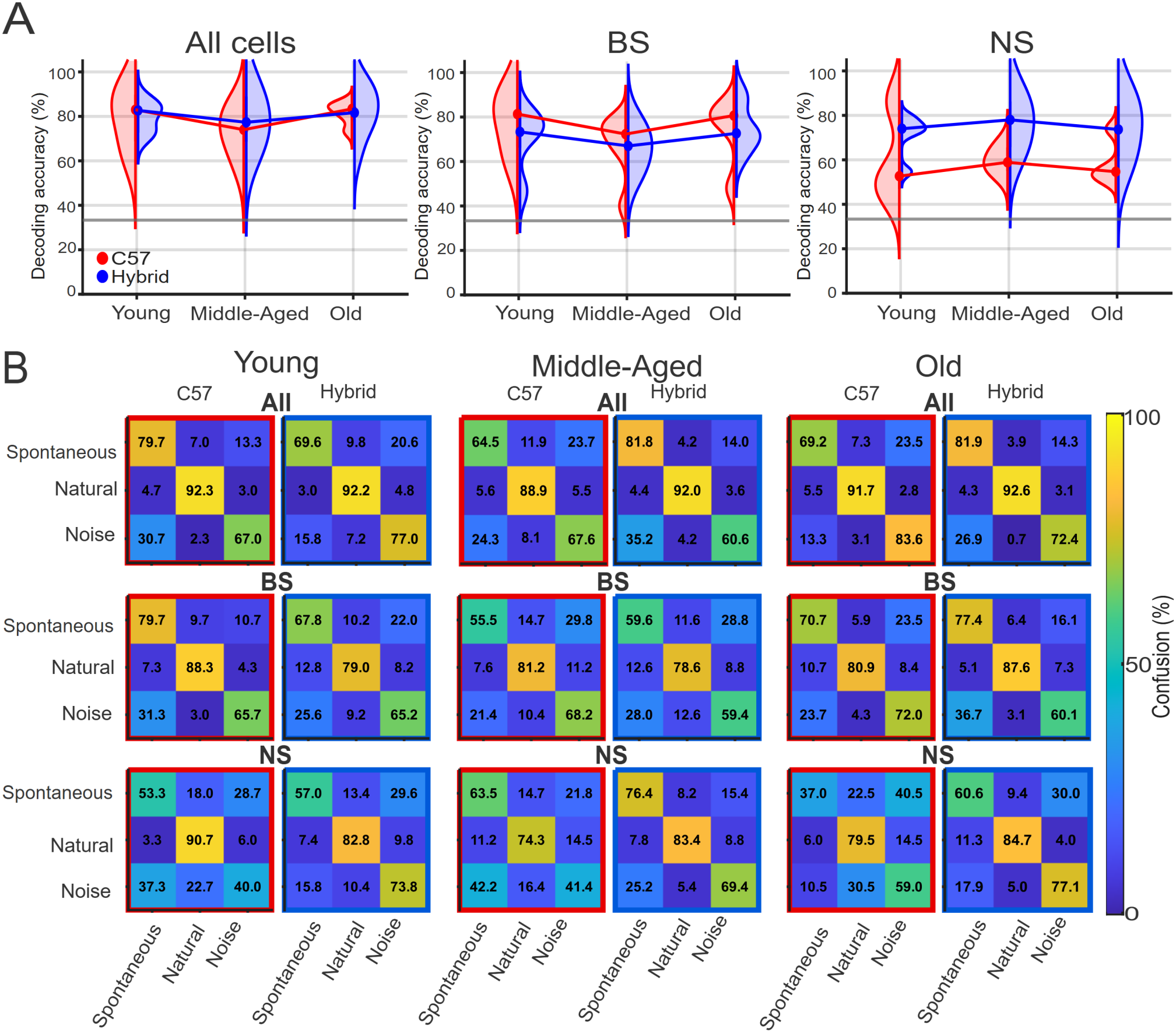
Population decoding of sound conditions across strains and age. **(A)** Decoding accuracy across young, middle-aged and old animals in C57 and Hybrid strains for All, BS and NS populations. Half-violin plots show the distribution of decoding accuracy across animals; central points indicate group medians and vertical lines indicate the IQR. n = 33 animals for the All population and n = 32 animals for BS and NS populations. The dashed line indicates the chance level (33.3%). No significant main effects of strain or age, or strain × age interactions, were detected in any population (two-way permutation ANOVA, 9,999 permutations; All: strain, F(1, 27) = 2.14, p = 0.160; age, F(2, 27) = 0.49, p = 0.609; strain × age, F(2, 27) = 0.36, p = 0.700; BS: strain, F(1, 26) = 2.62, p = 0.120; age, F(2, 26) = 0.99, p = 0.394; strain × age, F(2, 26) = 0.04, p = 0.966; NS: strain, F(1, 26) = 0.62, p = 0.443; age, F(2, 26) = 0.21, p = 0.808; strain × age, F(2, 26) = 0.76, p = 0.484, all Holm-adjusted p-values = 1.000). **(B)** Mean row-normalized confusion matrices for C57 and Hybrid animals across young, middle-aged and old age groups for All, BS and NS populations. Rows indicate the true stimulus class and columns the decoded class. Red and blue outlines denote C57 and Hybrid, respectively.

Decoding accuracy did not significantly correlate with age in either strain across the All, BS, or NS populations (Spearman’s rank correlation; C57: ρ = −0.475, p = 0.063 for All; ρ = −0.366, p = 0.179 for BS; ρ = −0.470, p = 0.077 for NS; Hybrid: ρ = −0.235, p = 0.365 for All; ρ = −0.296, p = 0.249 for BS; ρ = −0.007, p = 0.978 for NS). All correlations remained non-significant after Holm correction (all adjusted p ≥ 0.377). The pattern of errors was also similar across groups (**Fig. 5B**). Natural sound-evoked activity was classified most accurately, and most errors were confusions between spontaneous and noise-evoked activity, in every strain and age group.

A linear support vector machine and linear discriminant analysis gave similar results (**Fig. S2**); all three decoders performed significantly above chance (Template: 78.85 ± 9.57%, p = 2.82 × 10^−7^; SVM: 82.50 ± 8.74%, p = 2.81 × 10^−7^; LDA: 79.35 ± 10.37%, p = 2.81 × 10^−7^; mean ± SD; Wilcoxon signed-rank tests against chance). This significant above-chance performance was also observed in both C57 mice (all p ≤ 2.41 × 10^−4^) and Hybrid mice (all p = 1.60 × 10^−4^). These results indicate that auditory cortical population activity reliably discriminates sound conditions across the lifespan, in mice with and without hearing loss.

## Discussion

We recorded over 3,000 single units from the auditory cortex of awake mice across the lifespan, in the C57 strain that loses its hearing with age and in a hybrid strain that does not. We examined how aging and hearing loss alter the spiking of individual neurons and the discriminability of sound conditions from population activity. Three results are notable. First, aging increased the irregularity of spontaneous firing and reduced spike-timing precision in both strains. Second, a set of changes appeared only in C57 mice: a loss of responsive BS neurons, an age-related rise in spontaneous firing of BS neurons, more variable sound-evoked spike trains, weaker coupling between spontaneous and evoked firing in NS neurons and a sound-evoked change in CE that became negative with age. Third, despite these single-neuron changes, population activity discriminated silence, natural sound and noise accurately in every strain and age group.

### Reduced recruitment in aging and hyperactivity in hearing loss

The proportion of neurons driven by natural sounds fell with age in C57 mice for both cell classes and in Hybrid mice for NS neurons, and old C57 mice had fewer responsive BS and NS neurons than old Hybrid mice (**Fig. 1G**). A reduced fraction of responsive neurons is a consistent feature of aged auditory cortex in mice with preserved peripheral hearing (Shilling-Scrivo et al. 2021; Mittelstadt et al. 2026), suggesting that part of this loss reflects cortical aging itself. The additional loss in C57 mice indicates that degraded peripheral input further reduces the cortical response to complex sounds (Turner et al. 2005; de Villers-Sidani et al. 2010; Hughes et al. 2010).

Spontaneous activity moved in opposite directions in the two strains. BS and NS neurons in Hybrid mice became less active with age, whereas BS neurons in C57 mice became more active (**Fig. 2A**). Elevated spontaneous firing after cochlear damage may be explained by the compensatory increase in cortical excitability that follows loss of afferent input and that has been linked to weakened inhibition and increased neuronal gain (Kotak et al. 2005; Auerbach et al. 2014; Chambers et al. 2016; Resnik and Polley 2021). Our data extend these observations to a genetic model of age-related hearing loss recorded in awake animals.

Evoked responses of NS neurons declined with age in both strains. The shared decline in NS evoked firing fits a hearing-independent aging of inhibitory neurons. In contrast, the weaker coupling between spontaneous and evoked firing of NS neurons was not significant in young and old C57 mice (**Fig. 2B**), although slopes were not compared between strains, so this difference remains tentative.

### Aging degrades the temporal organization of spike trains

Aging increased the irregularity of spontaneous spike trains in both strains and reduced the precision with which neurons reproduced their spike timing across repeated natural sounds (**Fig. 3**). These changes are consistent with evidence that temporal processing declines with age independently of audiometric thresholds (Humes et al. 2012). Hearing loss added to them: sound-evoked spike trains became more variable with age only in C57 mice, and old C57’s NS neurons were less precise than old Hybrid’s NS neurons. This agrees with the finding that central gain after cochlear denervation restores rate-coded but not timing-dependent features of sound (Chambers et al. 2016). Because fast-spiking interneurons impose the millisecond-scale timing of cortical responses (Wehr and Zador 2003), a loss of their temporal precision is a plausible route from weakened inhibition to imprecise responses.

### Sound-evoked changes in spike-pattern entropy differ between strains

CE quantifies how fully a neuron uses the space of spike patterns available at its firing rate (Pryluk et al. 2019). The effect of natural sounds on CE diverged between strains with age. The median modulation was slightly positive in Hybrid’s neurons at every age, whereas in C57’s neurons it decreased with age and was negative in middle-aged and old mice (**Fig. 4**). The sign of the modulation within each strain was not tested, so the supported result is the difference between strains. Whether this shift preserves a degraded input through more stereotyped firing or simply reflects an impoverished input cannot be decided from these data.

### Population activity discriminates sound conditions across the lifespan

Population activity distinguished spontaneous, natural sound-evoked and noise-evoked conditions with high accuracy, and accuracy did not change with age or differ between strains (**Fig. 5**). This is consistent with the near-normal sound detection that survives extensive cochlear denervation (Chambers et al. 2016) and with the tolerance of population codes to degradation of their individual elements (Quian Quiroga and Panzeri 2009). The analysis suggests that rate-based information about the presence and broad category of sound is maintained, and it remains to be tested whether the discrimination of individual sounds, which depends more on spike timing, is equally robust.

### Aging versus hearing loss

Comparing the two strains separates two components of cortical aging. Changes present in both strains, and therefore attributable to aging with preserved hearing, included the loss of responsive NS neurons, the decline of NS evoked firing, more irregular spontaneous firing and reduced spike-timing precision of NS neurons. Changes seen only with hearing loss included the loss of responsive BS neurons, the rise in spontaneous firing of BS neurons, more variable sound-evoked spike trains and a negative sound-driven change in CE. This division agrees with the view that the loss of parvalbumin neurons and of intracortical connectivity arises largely independently of cochlear damage (Rogalla and Hildebrandt 2020; Maximov and Kanold 2025; Xu et al. 2025), and adds that hearing loss superimposes hyperexcitability and altered evoked spike patterns.

### Limitations of the study

BS and NS classes were defined from spike waveforms, which identify excitatory and fast-spiking inhibitory neurons only approximately (Lee et al. 2021). The Hybrid strain carries one CBA/Ca genome, so strain differences may reflect CBA/Ca-derived alleles other than the complemented Cdh23 locus. The design was cross-sectional and mice listened passively, so the relationship between these changes and perception remains to be determined.

## Conclusion

Aging makes the spiking of auditory cortical neurons less regular and less precise. Age-related hearing loss adds a loss of responsive neurons, hyperexcitability of BS neurons and altered sound-evoked spike patterns. Population activity nevertheless continues to discriminate sound conditions accurately. Age-related hearing loss may therefore affect cortical processing mainly through the recruitment of neurons and the timing and structure of their spike trains.

## Funding

This work was supported by the Biotechnology and Biological Sciences Research Council (grant number BB/M00905X/1) and the Medical Research Council (grant number MR/Y004051/1) to S.S.

## Acknowledgments

We thank Daniel Lyngholm for contributing to the recordings and for characterizing the hearing thresholds of the mouse lines used here, and members of the Sakata laboratory for discussion.

## Author contributions

Sarah Fouda (Conceptualization, Data curation, Formal analysis, Investigation, Methodology, Resources, Software, Validation, Visualization, Writing – original draft, Writing – review & editing) and Shuzo Sakata (Conceptualization, Funding acquisition, Project administration, Supervision, Writing – original draft, Writing – review & editing).

## Conflict of interest statement

The authors declare no competing interests.

## Data and code availability

Spike-sorted data tables and the MATLAB and R code used for all analyses will be deposited in a public repository upon publication.

## Notes

### Competing Interest Statement

The authors have declared no competing interest.

## References

Anderson MJ. 2001. A new method for non-parametric multivariate analysis of variance. Austral Ecol. 26:32–46. 10.1111/j.1442-9993.2001.01070.pp.x

Auerbach BD, Rodrigues PV, Salvi RJ. 2014. Central gain control in tinnitus and hyperacusis. Front Neurol. 5:206. 10.3389/fneur.2014.00206

Bowen Z, Winkowski DE, Kanold PO. 2020. Functional organization of mouse primary auditory cortex in adult C57BL/6 and F1 (CBAxC57) mice. Sci Rep. 10:10905. 10.1038/s41598-020-67819-4

Caspary DM, Ling L, Turner JG, Hughes LF. 2008. Inhibitory neurotransmission, plasticity and aging in the mammalian central auditory system. J Exp Biol. 211:1781–1791. 10.1242/jeb.013581

Caspary DM, Hughes LF, Ling LL. 2013. Age-related GABAA receptor changes in rat auditory cortex. Neurobiol Aging. 34:1486–1496.10.1016/j.neurobiolaging.2012.11.009

Chambers AR, Resnik J, Yuan Y, Whitton JP, Edge AS, et al.2016. Central gain restores auditory processing followingnear-complete cochlear denervation. Neuron. 89:867– 879. 10.1016/j.neuron.2015.12.041

de Ruyter van Steveninck RR, Lewen GD, Strong SP, Koberle R, Bialek W. 1997. Reproducibility and variability in neural spike trains. Science. 275:1805–1808. 10.1126/science.275.5307.1805

de Villers-Sidani E, Alzghoul L, Zhou X, Simpson KL, Lin RCS, et al. 2010. Recovery of functional and structural age-related changes in the rat primary auditory cortex with operant training. Proc Natl Acad Sci U S A.107:13900–13905.10.1073/pnas.1007885107

Flurkey K, Currer JM, Harrison DE. 2007. Mouse models in aging research. In: Fox JG, Barthold SW, Davisson MT, Newcomer CE, Quimby FW, et al., editors. The mouse in biomedical research. 2nd ed. Vol. III, Normative biology, husbandry, and models. Academic Press. p. 637–672. 10.1016/B978-012369454-6/50074-1

Frisina RD, Singh A, Bak M, Bozorg S, Seth R, et al. 2011. F1 (CBA×C57) mice show superior hearing in old age relative to their parental strains: hybrid vigor or a new animal model for “golden ears”? Neurobiol Aging.32:1716–1724.10.1016/j.neurobiolaging.2009.09.009

Holm S. 1979. A simple sequentially rejective multiple testprocedure. Scand J Stat. 6:65–70.

Hughes LF, Turner JG, Parrish JL, Caspary DM. 2010. Processing of broadband stimuli across A1 layers in young and aged rats. Hear Res. 264:79–85. 10.1016/j.heares.2009.09.005

Humes LE, Dubno JR, Gordon-Salant S, Lister JJ, Cacace AT, et al. 2012. Central presbycusis: a review and evaluation of the evidence. J Am Acad Audiol. 23:635–666. 10.3766/jaaa.23.8.5

Kotak VC, Fujisawa S, Lee FA, Karthikeyan O, Aoki C, et al.2005. Hearing loss raises excitability in the auditory cortex. J Neurosci. 25:3908–3918. 10.1523/JNEUROSCI.5169-04.2005

Kujawa SG, Liberman MC. 2009. Adding insult to injury: cochlear nerve degeneration after “temporary” noise-induced hearing loss. J Neurosci. 29:14077–14085. 10.1523/JNEUROSCI.2845-09.2009

Lee EK, Balasubramanian H, Tsolias A, Anakwe SU, Medalla M, et al. 2021. Non-linear dimensionality reduction on extracellular waveforms reveals cell type diversity in premotor cortex. eLife. 10:e67490. 10.7554/eLife.67490

Lin FR, Niparko JK, Ferrucci L. 2011. Hearing loss prevalence in the United States. Arch Intern Med. 171:1851–1852.10.1001/archinternmed.2011.506

Ling LL, Hughes LF, Caspary DM. 2005. Age-related loss of the GABA synthetic enzyme glutamic acid decarboxylase in rat primary auditory cortex. Neuroscience. 132:1103– 1113. 10.1016/j.neuroscience.2004.12.043

Lyngholm D, Sakata S. 2019. Cre-dependent optogenetic transgenic mice without early age-related hearing loss. Front Aging Neurosci. 11:29. 10.3389/fnagi.2019.00029

Martin del Campo HN, Measor KR, Razak KA. 2012.Parvalbumin immunoreactivity in the auditory cortex of a mouse model of presbycusis. Hear Res. 294:31–39. 10.1016/j.heares.2012.08.017

Maximov K, Kanold PO. 2025. Aging reduces excitatory bandwidth, alters spectral tuning curve diversity, and reduces sideband inhibition in L2/3 of primary auditory cortex. bioRxiv. 10.1101/2025.04.02.646797, preprint: not peer reviewed.

Mittelstadt JK, Shilling-Scrivo KV, Kanold PO. 2026. Aging in the primary auditory cortex. J Assoc Res Otolaryngol. 27:37–56. 10.1007/s10162-025-01013-z

Ouellet L, de Villers-Sidani E. 2014. Trajectory of the mainGABAergic interneuron populations from earlydevelopment to old age in the rat primary auditory cortex.Front Neuroanat. 8:40. 10.3389/fnana.2014.00040

Pryluk R, Kfir Y, Gelbard-Sagiv H, Fried I, Paz R. 2019. A tradeoff in the neural code across regions and species. Cell. 176:597–609.e18.10.1016/j.cell.2018.12.032

Quian Quiroga R, Panzeri S. 2009. Extracting information from neuronal populations: information theory and decoding approaches. Nat Rev Neurosci. 10:173–185. 10.1038/nrn2578

Resnik J, Polley DB. 2021. Cochlear neural degeneration disrupts hearing in background noise by increasing auditory cortex internal noise. Neuron. 109:984–996.e4. 10.1016/j.neuron.2021.01.015

Rieke F, Warland D, de Ruyter van Steveninck R, Bialek W.1999. Spikes: exploring the neural code. MIT Press.

Rogalla MM, Hildebrandt KJ. 2020. Aging but not age-related hearing loss dominates the decrease of parvalbumin immunoreactivity in the primary auditory cortex of mice. eNeuro. 7:ENEURO.0511-19.2020.10.1523/ENEURO.0511-19.2020

Schmitzer-Torbert N, Jackson J, Henze D, Harris K, Redish AD. 2005. Quantitative measures of cluster quality foruse in extracellular recordings. Neuroscience. 131:1–11. 10.1016/j.neuroscience.2004.09.066

Schreiber S, Fellous JM, Whitmer D, Tiesinga P, Sejnowski TJ. 2003. A new correlation-based measure of spike timing reliability. Neurocomputing. 52–54:925–931. 10.1016/S0925-2312(02)00838-X

Sergeyenko Y, Lall K, Liberman MC, Kujawa SG. 2013. Age-related cochlear synaptopathy: an early-onset contributor to auditory functional decline. J Neurosci. 33:13686–3694. 10.1523/JNEUROSCI.1783-13.2013

Shilling-Scrivo K, Mittelstadt J, Kanold PO. 2021. Altered response dynamics and increased population correlation to tonal stimuli embedded in noise in aging auditory cortex. J Neurosci. 41:9650–9668. 10.1523/JNEUROSCI.0839-21.2021

Turner JG, Hughes LF, Caspary DM. 2005. Affects of aging on receptive fields in rat primary auditory cortex layer V neurons. J Neurophysiol. 94:2738–2747. 10.1152/jn.00362.2005

Wehr M, Zador AM. 2003. Balanced inhibition underlies tuning and sharpens spike timing in auditory cortex. Nature. 426:442–446.10.1038/nature02116

Willott JF, Aitkin LM, McFadden SL. 1993. Plasticity of auditory cortex associated with sensorineural hearing loss in adult C57BL/6J mice. J Comp Neurol. 329:402–411. 10.1002/cne.903290310

Wu PZ, Liberman LD, Bennett K, de Gruttola V, O’Malley JT, et al. 2019. Primary neural degeneration in the human cochlea: evidence for hidden hearing loss in the aging ear. Neuroscience. 407:8–20. 10.1016/j.neuroscience.2018.07.053

Xu Z, Xue B, Kao JPY, Kanold PO. 2025. Sex-specific age-related changes in excitatory and inhibitory intracortical circuits in mouse primary auditory cortex. eNeuro.12:ENEURO.0378-24.2024.10.1523/ENEURO.0378-24.2024

Xue B, Meng X, Kao JPY, Kanold PO. 2023. Age-related changes in excitatory and inhibitory intra-cortical circuits in auditory cortex of C57Bl/6 mice. Hear Res.429:108685.10.1016/j.heares.2022.108685

