## Supplementary Information for "Distinct effects of aging and hearing loss on mouse auditory cortex"

*Supplementary Figures S1 and S2.*

### Supplementary figures

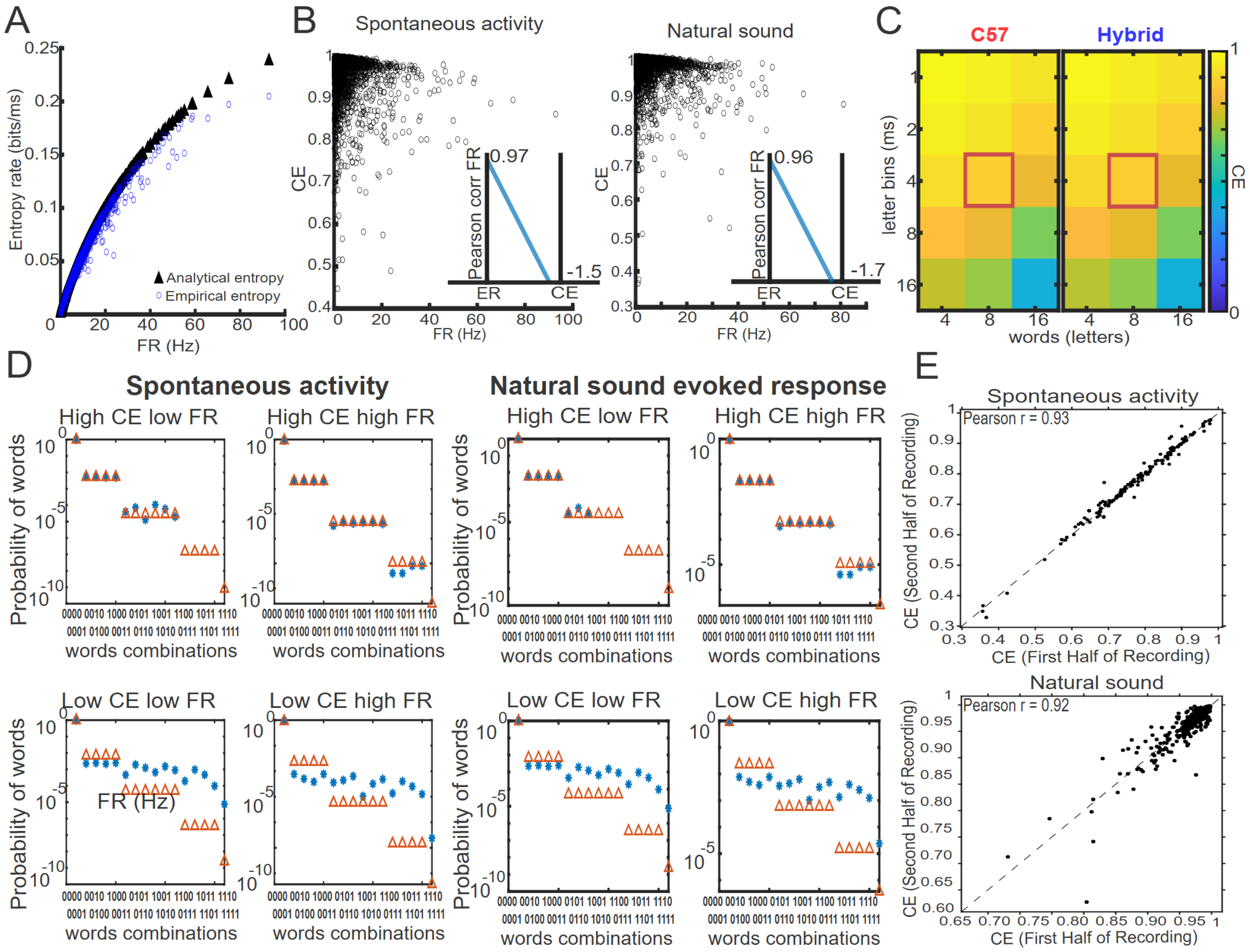


**Figure S1. Contrast entropy calculation during spontaneous and natural sound-evoked activity.**

**(A)** Entropy rate (ER) as a function of firing rate (FR) during spontaneous activity. ER increased steeply with FR (Pearson's r = 0.97 during spontaneous activity and 0.96 during natural sound stimulation).

**(B)** Contrast entropy (CE) (Pryluk et al. 2019) as a function of FR during spontaneous activity (left) and natural sound stimulation (right). Conversion from ER to CE removes the positive dependence on FR (Pearson's r = −0.15 during spontaneous activity and −0.17 during natural sound stimulation).

**(C)** CE across word lengths and letter bin widths for C57 (red) and Hybrid (blue) neurons. Red boxes indicate the parameters selected for subsequent analyses (4-ms letter bins and 8 letters per word).

**(D)** Agreement between analytical and observed word probabilities during spontaneous activity (left) and natural sound presentation (right). Red triangles represent analytical entropy-derived probabilities; blue asterisks represent word probabilities computed from recorded neurons. The same neurons are shown during spontaneous activity (left) and natural sound presentation (right) across different CE/FR regimes. For each condition, from left to right: CE = 0.99, FR = 1.79; CE = 0.99, FR = 10.63; CE = 0.70, FR = 23.73; and CE = 0.86, FR = 23.74.

**(E)** Comparison of CE estimates between the first and second halves of recordings during spontaneous activity (top) and natural sound stimulation (bottom). Each point represents an individual recording.

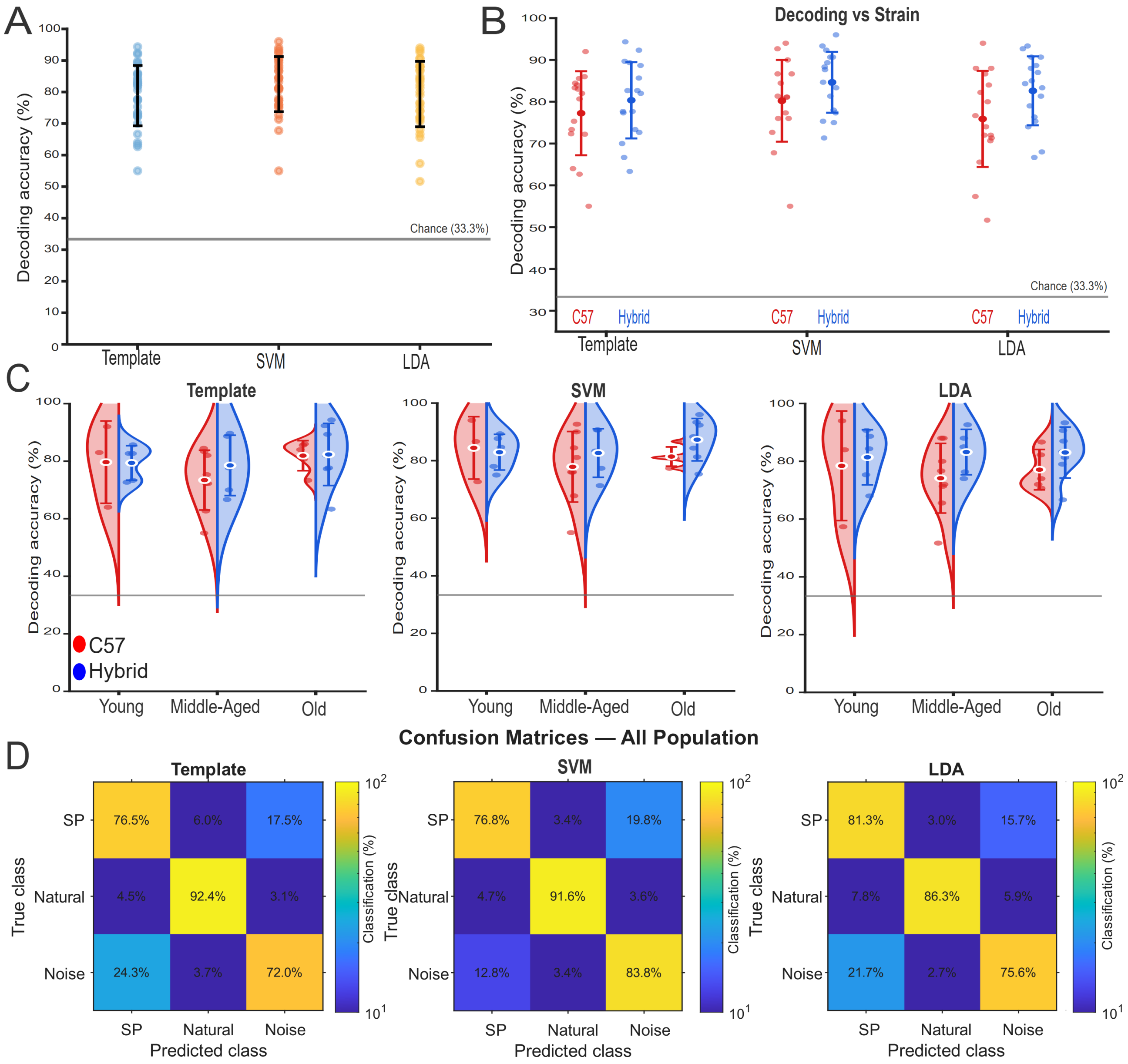


**Figure S2. Decoding performance and confusion matrices across decoders.**

**(A)** Comparison of decoding accuracy across Template, SVM, and LDA decoders. Each point represents one observation; horizontal bars indicate mean ± SD. The dashed horizontal line indicates chance-level performance (33.3%). Decoding accuracy was significantly above chance for all three decoders (one-sided Wilcoxon signed-rank tests: Template, $p = 2.82 \times 10^{-7}$; SVM, $p = 2.81 \times 10^{-7}$; LDA, $p = 2.81 \times 10^{-7}$). Linear SVM achieved the highest mean decoding accuracy (82.50 ± 8.74%) and significantly outperformed both the Template decoder (78.85 ± 9.57%; paired t-test, Holm-adjusted $p = 0.0056$) and LDA (79.35 ± 10.37%; Holm-adjusted $p = 0.0010$). Template and LDA did not differ significantly (Holm-adjusted $p = 0.6662$).

**(B)** Decoding accuracy across strains. Decoding accuracy (%) is shown separately for C57 and Hybrid animals for each decoder. Each point represents one observation; bars indicate mean ± SD. No significant strain effect was detected for Template, Linear SVM, or LDA decoding (Template, $F(1, 27) = 0.244$, $p = 0.611$; Linear SVM, $F(1, 27) = 0.896$, $p = 0.325$; LDA, $F(1, 27) = 2.405$, $p = 0.151$). Accuracy was numerically higher in Hybrid mice for all three decoders (Template: 80.35 ± 9.13% vs 77.24 ± 10.05%; Linear SVM: 84.65 ± 7.28% vs 80.22 ± 9.79%; LDA: 82.61 ± 8.25% vs 75.88 ± 11.49%); however, no significant strain effect was detected after Holm correction (Template, Holm-adjusted $p = 0.651$; Linear SVM, Holm-adjusted $p = 0.651$; LDA, Holm-adjusted $p = 0.454$).

**(C)** Decoding accuracy by age and strain. Decoding accuracy is shown for young, middle-aged, and old animals separately for C57 and Hybrid strains for Template, Linear SVM, and LDA decoding. Violin distributions show the observed data, with individual observations overlaid and mean ± SD indicated. No significant age × strain interaction was detected for any decoder (two-way ANOVA; Template, $F(2, 27) = 0.247$, $p = 0.783$; Linear SVM, $F(2, 27) = 0.439$, $p = 0.649$; LDA, $F(2, 27) = 0.199$, $p = 0.820$).

**(D)** Confusion matrices for the All population. Mean confusion matrices are shown for Template, SVM, and LDA decoders. Rows indicate the true class and columns indicate the predicted class. Values represent the percentage of trials classified into each category. Natural sounds showed the highest classification accuracy across all three decoding methods. n = 33 mice.
